# Honey bees (*Apis mellifera*) exhibit seasonal variation in their tolerance to viral infection

**DOI:** 10.1101/2025.07.11.664207

**Authors:** Alexandria N. Payne, Ashley L. St. Clair, Gyan P. Harwood, Vincent Prayugo, Lincoln N. Taylor, Madeleine Shapiro, Adam G. Dolezal

## Abstract

Seasonal variation strongly influences honey bee colony dynamics, leading to time-dependent changes in behavioral and physiological phenotypes. However, the extent to which seasonal fluctuations affect the susceptibility and tolerance of honey bees to viral infection remains largely unexplored. To address this, we conducted a longitudinal study in which adult honey bee workers were collected monthly from research colonies and experimentally infected with Israeli acute paralysis virus (IAPV) over the course of a year. Our results showed significant seasonal variation in the mortality and IAPV load of inoculated bees, with bees challenged during the pre-overwintering period (i.e., fall) exhibiting the highest susceptibility and lowest tolerance to IAPV infection compared to bees challenged in spring, summer, or winter. To investigate factors underlying these seasonal differences, we conducted nutrition-based studies that determined: 1) the variation in lipid content of colonies throughout the year and its potential link to our observed trends in IAPV tolerance, and 2) the impact of seasonally collected pollen on the survivorship of IAPV-challenged bees. Our findings support that seasonal changes in honey bee physiology and nutritional status play key roles in influencing honey bee viral tolerance. We conclude that honey bee colonies are particularly vulnerable to viral infection during the pre-overwintering period, most likely as a result of reduced tolerance to pathogen stress when transitioning from a summer to winter worker population. We further hypothesize that this period of increased vulnerability to viral infection, in correlation with other disease factors such as *Varroa* mite abundance and available forage, likely contributes to the relatively high overwintering losses experienced by beekeepers. Given the recent reports of severe colony losses attributed to honey bee viruses, understanding the relationship between seasonality and viral tolerance in honey bees is crucial for better informing management strategies and improving overwintering success.

**AUTHOR SUMMARY:** We explored how seasonal changes affect the ability of honey bees to withstand viral infections. Previous research has shown that there are physical and behavioral differences between summer and winter bees, but it’s unclear how these seasonal differences affect a honey bee’s ability to withstand viral infection. To investigate this, we collected honey bee workers monthly and infected them with Israeli acute paralysis virus (IAPV) over the course of a year. Our results showed that honey bees were most vulnerable to IAPV in the fall (i.e., prior to overwintering), as they showed the highest mortality rates and lowest viral tolerance, based on their IAPV loads, during this time. By following up with nutrition-based studies, we found that seasonal changes in bee nutrition in part explained the seasonal differences we observed in honey bee virus tolerance. Overall, our findings suggest that bees are less tolerant to viral infection during the pre-overwintering period when colonies transition from a summer to a winter worker population.

This vulnerable period may help explain the high rates of colony losses experienced by beekeepers nationwide and demonstrates the importance of developing seasonally-dependent disease management strategies.

## 1. INTRODUCTION

Seasonal variation plays a critical role in shaping the disease dynamics of many host-pathogen systems (1–4). This can be illustrated with conceptual models such as the disease triangle (5), which considers temporal effects (e.g., seasonality) when predicting epidemiological outcomes between a host, an infectious agent, and the environment (6).

Seasonality has been shown to affect the relationship between honey bees (*Apis mellifera*) and their viral pathogens (7–9), with viral abundance being closely correlated to seasonal trends in *Varroa* mite (*Varroa destructor*) populations (reviewed in: Genersch and Aubert, 2010; Traynor et al., 2020) due to these mites being highly effective vectors of many honey bee-infecting viruses (12,13). *Varroa* populations, along with viral abundance, typically peak in colonies during the mid-summer to fall period (10,11), often increasing incidences of virus transmission and risk of exposure. Elevated levels of this *Varroa*-virus complex during the transitional period in which honey bee colonies prepare to overwinter (i.e., the pre-overwintering period) has been identified as a primary driver behind the unsustainable rates of overwintering colony losses experienced by beekeepers (14–18) and may be a cause of the recent mass colony losses observed in 2025 (19).

Increased *Varroa* abundance alone, however, does not fully explain overwintering outcomes, as seasonally-dependent changes in host physiology and environmental conditions also critically influence a colony’s pathogen tolerance and overall survival, with tolerance defined as a host’s ability to mitigate infection-related fitness costs without directly affecting pathogen load (20). For example, seasonal fluctuations in food availability, particularly in forage resources (21,22), have been shown to impact honey bee health and colony survival (23,24).

These seasonal fluctuations can lead to poor host nutrition, which can negatively affect honey bee physiology, immune function, and pathogen tolerance at both the individual and colony level (25,26). While geographically dependent, the pre-overwintering period typically corresponds with relatively reduced forage, often resulting in the decreased nutritional health of honey bee colonies in combination with increased *Varroa* and virus levels during this transitional time.

Additionally, intrinsic host factors, such as an individual’s physiology, are also known to influence host tolerance and susceptibility, with susceptibility defined as the probability that an infection will occur (6,27,28). Given that honey bees exhibit seasonally-dependent phenotypes, with significant physiological and behavioral differences between summer and winter worker populations (29–37), it stands to reason that honey bee colonies would also exhibit seasonality in their tolerance to viral infection throughout their yearly cycle. However, this is difficult to determine and disentangle from the aforementioned pathogen-and environment-related factors (e.g., *Varroa*/viral abundance and available forage) that are influenced by seasonality and known to shape honey bee tolerance.

To better understand how seasonality influences honey bee tolerance to pathogen infection, we conducted a longitudinal study that challenged honey bees with viral infection over the course of a year by means of monthly dose-response trials. Cohorts of adult worker bees were experimentally infected with Israeli acute paralysis virus (IAPV), which was selected due to its controllable inoculum potential (38), distinct and rapid mortality symptoms (39), and its previous use as a model honey bee pathogen (40–44). We hypothesized that honey bees exhibit seasonal variation in their tolerance to viral infections, even when extrinsic disease factors, such as *Varroa*, are controlled for. Then, to investigate potential contributing factors related to our findings, we performed two nutrition-associated studies where we: 1) took longitudinal measurements of colony-level lipid content as an indicator of host nutritional status, and 2) fed IAPV-inoculated bees with season-specific bee-collected pollen diets to assess the influence of seasonal nutrition on pathogen response.

Overall, we found that honey bees experience seasonal differences in their tolerance to viral infection, with bees challenged in the pre-overwintering period (i.e., fall) showing the lowest tolerance, most likely as a result of seasonal differences in available forage and host nutritional status. These findings further our understanding of the role that seasonality plays in honey bee tolerance, which is critical for developing targeted strategies to mitigate disease risks and better management strategies to improve overwintering colony survival.

## 2. RESULTS

### 2.1 IAPV-induced mortality varies based on month of inoculation

A longitudinal viral dose-response assay (hereafter referred to as the longitudinal tolerance experiment) revealed that honey bee mortality varied significantly depending on the month in which IAPV inoculation occurred. Mortality rates of experimentally infected honey bees at 48 hours post-inoculation (hpi) varied significantly by trial month (d.f. = 11, χ^2^ = 183.88, *p* < 0.0001), IAPV treatment (d.f. = 7, χ^2^ = 445.64, *p* < 0.0001), and by the interaction between trial month and IAPV treatment (d.f. = 77, χ^2^ = 172.60, *p* < 0.0001), where IAPV treatment consisted of a handling control group, an injection control group, and six IAPV inoculum treatments (high dose, 10^-2^, 10^-4^, 10^-6^, 10^-8^, and 10^-10^), as described in section 4.2.

Cumulative bee mortality at 48 hpi (Figure 1A) for each trial month (February 2021-January 2022) and each IAPV treatment showed that only the two most concentrated IAPV inoculum treatments (i.e., the high dose and 10^-2^ dilution) resulted in over 50% mortality across all trial months. Chi-square analyses compared the cumulative mortality at 48 hpi of the injection control group against each other IAPV treatment within each trial month (Table S1) and showed that the high dose and 10^-2^ treatments were the only groups that significantly differed from their respective injection controls across all trial months, aside from the high dose group in September. The other IAPV treatments varied in their survivorship over time, particularly the 10^-4^ dose (Figure 1B). Interestingly, the 10^-4^ IAPV treatment resulted in 100% mortality of inoculated bees in only the month of August. In fact, August had the highest overall mortality, with the high dose and five dilution treatments resulting in >75% mortality and significantly differing from their respective injection controls. While September also showed a similarly high level of mortality across the IAPV inoculum treatments, it also displayed an abnormally high level of mortality for both the injection and handling control groups.

**Figure 1:**
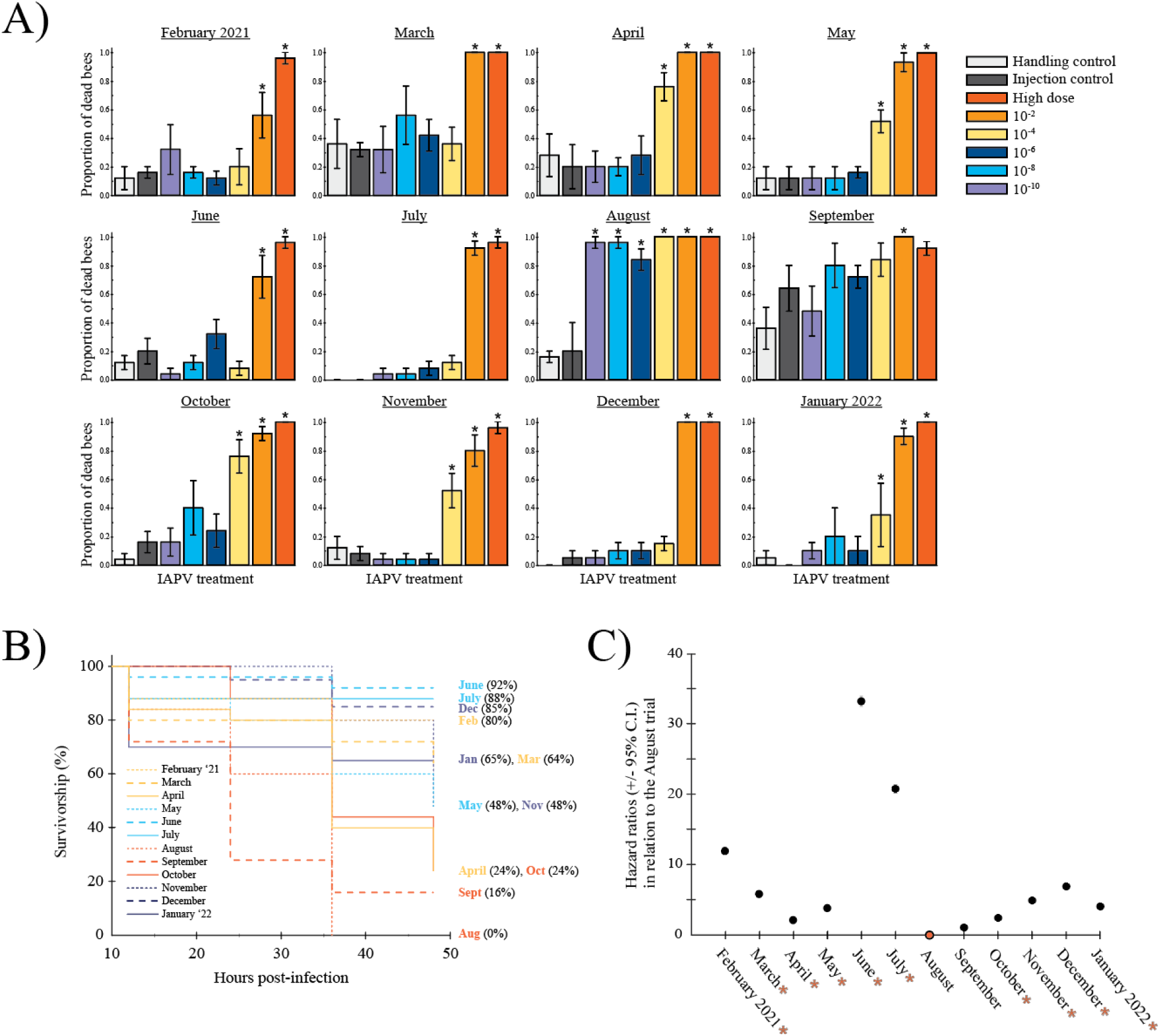
*IAPV-induced mortality was dependent on the month in which inoculation occurred.* **A)** Cumulative mortality at 48 hpi for each IAPV treatment group (n=8) across all trial months (n=12). The different colored bars represent the high dose of IAPV (∼9.86 x 10^3^ g.e. of IAPV within 100 ng of purified total RNA), its five dilutions (10^-2^, 10^-4^, 10^-6^, 10^-8^, 10^-10^), and two negative controls (handling control and injection control). Within each month, asterisks above bars indicate the IAPV treatments that significantly differed from the injection control (dark gray bars), based on Chi-square tests (*p* < 0.05). **B)** Survivorship curves for the 10^-4^ treatment over 48 hpi, excluding mortality prior to 12 hpi due to cage mishandling and not IAPV-induced mortality. Colored lines depicting each month are grouped by season (yellow= spring, blue= summer, orange= fall, purple= winter). **C)** Estimated hazard ratios (± 95% C.I.) of the 10^-4^ IAPV treatment group at 48 hpi, using the August trial (i.e., the trial month with the lowest survival at 48 hpi) as the reference comparison. Asterisks on the months along the x-axis indicate significant comparisons to the August reference comparison (Cox proportional hazards model, *p* < 0.05).

Survivorship analyses for the 10^-4^ IAPV dose (i.e., the dose with the highest variability in response across the year) confirmed the significant influence of trial month on the likelihood of bee death following IAPV inoculation (Figure 1C; d.f. = 14.22, χ^2^ = 119.9, *p* < 0.0001). Bees inoculated with the 10^-4^ dose in August had a significantly higher risk of mortality compared to bees inoculated in other months (Table S2), aside from September, which had a similar level of risk (Cox proportional hazards: HR= 1.07, *p* = 0.82). The greatest discrepancy in mortality risk was observed between bees inoculated in August vs those inoculated in the summer months of June and July. Bees inoculated with IAPV in August were more than 30 times more likely to die compared to bees inoculated in June, and more than 20 times more likely to die compared to bees inoculated in July.

### 2.1 Honey bee tolerance to IAPV infection is seasonally dependent

We quantified the IAPV load of a subset of honey bee samples collected during the longitudinal tolerance experiment. The IAPV load of bees that had died as a result of injection with the 10^-2^ IAPV treatment group was quantified and then grouped by season (spring= February and March, summer= May and June, fall= August and September, and winter= November and December).

We determined that honey bee tolerance to IAPV infection varied by season (Figure 2; Kruskal-Wallis rank sum test: d.f. = 3, χ^2^ = 28.53, *p* < 0.0001). Across all seasons, the mean IAPV load exceeded the threshold for what is considered a high-level infection (>10^4^ genome equivalents/ 20 ng of RNA) as defined in previous studies (45,46). Pairwise comparisons using Wilcoxon rank sum tests revealed significantly lower mean IAPV quantity, and greater degree of variation between samples, in fall-inoculated bees compared to bees inoculated in the spring (*p* < 0.0001), summer (*p* = 0.0176), or winter (*p* = 0.0141). Significant differences were also observed between spring and summer bees (*p* = 0.0431) and between spring and winter bees (*p* = 0.0002).

**Figure 2:**
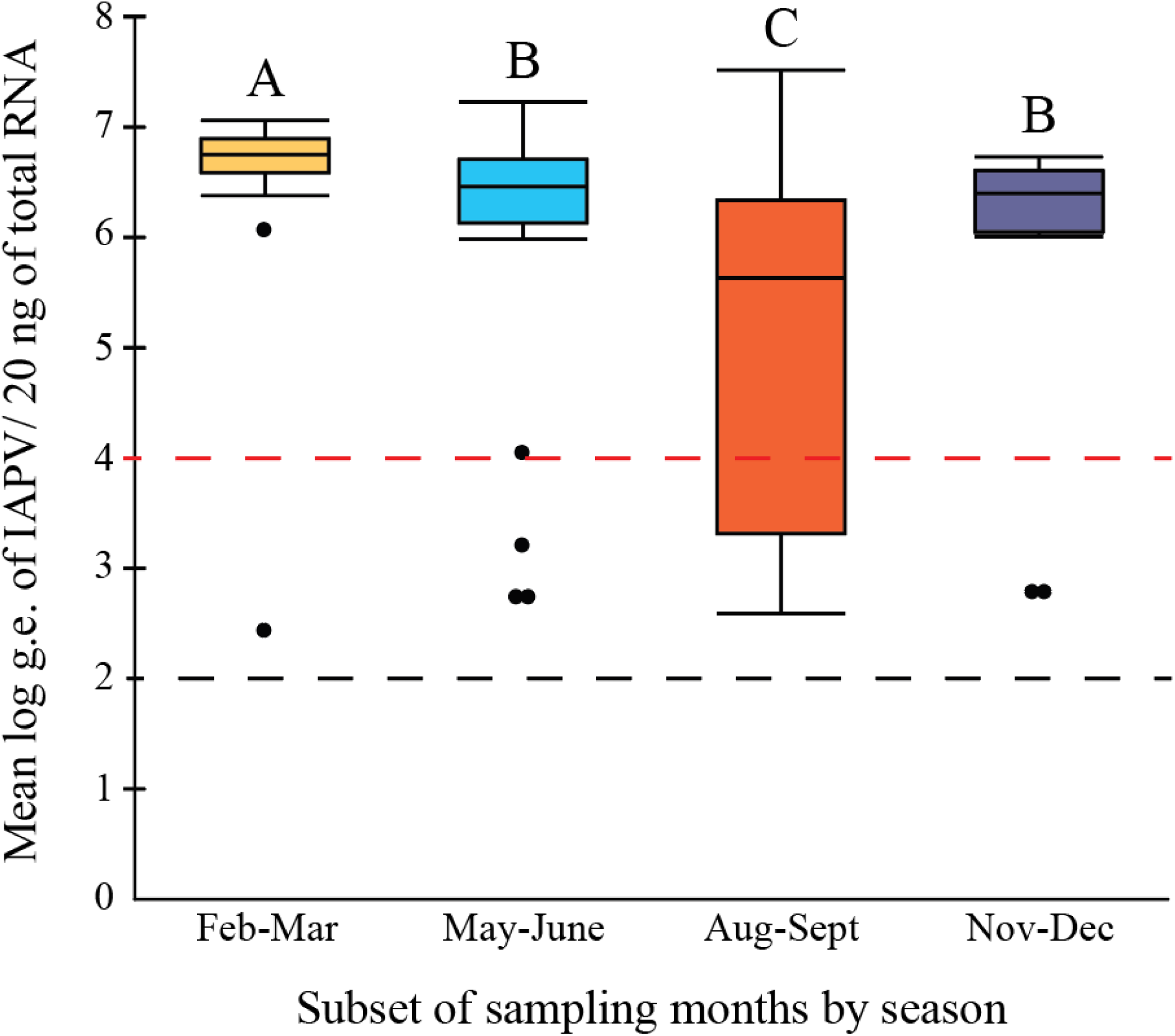
*Honey bee tolerance to IAPV infection was lowest during the pre-overwintering period (i.e., August-September).* We quantified the IAPV loads of individual, whole-bodied bees from the 10^-2^ IAPV treatment group that had died within 12-48 hpi during the longitudinal tolerance experiment. Bees were grouped by season (yellow= spring, blue= summer, orange= fall, purple= winter), with each season including samples spanning two trial months: February & March (spring; n=22), May & June (summer; n=24), August & September (fall; n=24), and November & December (winter; n=24). Boxes represent the mean (± s.e.) log genome equivalents (g.e.) of IAPV/ 20 ng of total extracted RNA. The black dashed line indicates the IAPV detection threshold (200 g.e. of IAPV/ reaction), and the red dashed line indicates the high-level IAPV infection threshold (>10^4^ g.e./ 20 ng of total RNA). Significant differences (α = 0.05) are indicated by differing letters (Kruskal-Wallis Test, Steel-Dwass test, *p* < 0.05).

However, summer and winter bees did not significantly differ from one another.

To confirm that observed IAPV loads resulted from experimental inoculations rather than pre-existing infections, we quantified the background IAPV levels of colonies used in the longitudinal tolerance experiment (Table S3) for the months of June (n=5 colonies), August (n=5), and December (n=4) in 2021. No IAPV was detected in any of the samples, ensuring that the observed differences in IAPV quantity between treatments were due to experimental infection. Additionally, *Varroa* populations within colonies remained on average below the 3% economic threshold (47) throughout the experiment (Figure S1), supporting that the results were not influenced by *Varroa* abundance. Finally, we quantified the background levels of deformed wing virus (DWV), black queen cell virus (BQCV), and sacbrood virus (SBV). Although above the limits of detection (Table S3), DWV and SBV quantities were consistently below thresholds associated with symptomatic infection (48,49). However, BQCV loads were elevated in two colonies in June and in one colony in August.

### 2.1 Colony-level lipid stores varied seasonally

To determine if seasonality in honey bee nutritional state relates to our observed seasonal variation in IAPV tolerance, we measured colony-level lipid content (i.e., fat stores) in parallel with the longitudinal tolerance experiment using the same research colonies. The mean percent bee lipid content significantly varied by trial month (Figure 3A; F_11,41_ = 3.15, *p* = 0.0036). Bees sampled in November had the lowest mean percent lipid (1.9%), significantly differing from bees with the highest mean percent lipid sampled in June (Tukey HSD: *p* = 0.0074), September (Tukey HSD: *p* = 0.0063), and January 2022 (Tukey HSD: *p* = 0.029), which all had approximately 4% lipid content.

**Figure 3:**
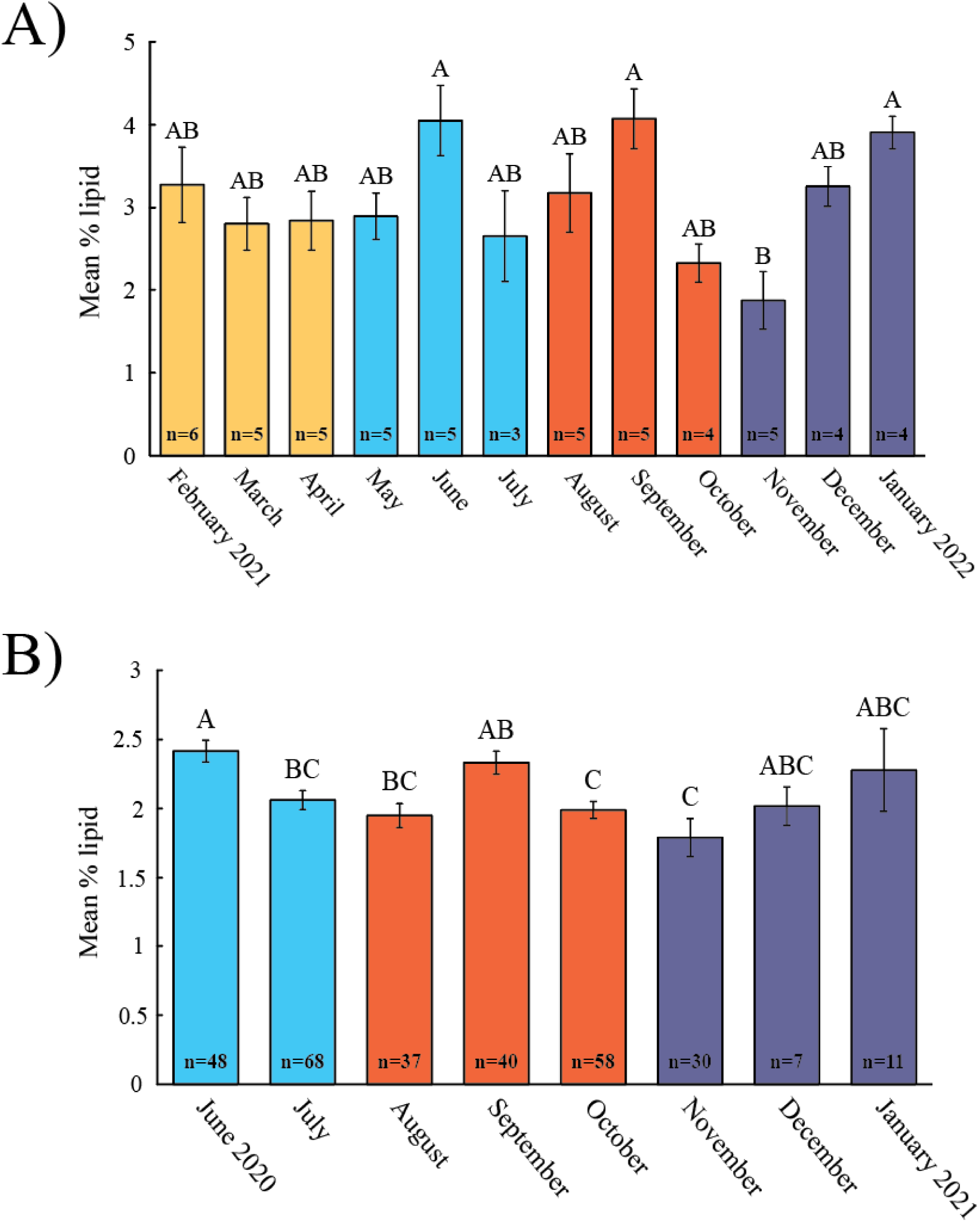
*Colony-level nutritional status varies seasonally.* **A)** Nutritional status of the honey bee colonies used in the longitudinal tolerance experiment, as estimated by their mean % lipid content (i.e. fat stores). **B)** The nutritional status of a larger pool of colonies sampled from the previous year (June 2020-January 2021) to establish a baseline of seasonal variation in honey bee colony nutrition and confirm trends from the smaller-scale, longitudinal tolerance experiment. In both panels, bars represent the mean (± s.e) percent lipid of all colonies by month, with the number of colonies sampled from each month depicted at each bar’s base. The color of each bar is based off of its seasonal grouping (yellow= spring, blue= summer, orange= fall, purple= winter). Significant differences are indicated by differing letters above each bar (α = 0.05).

To confirm these findings, we conducted a second longitudinal study using a much larger sample size of 80 colonies from which honey bees had previously been collected from June 2020-January 2021. This study resulted in a similar trend as our 2021-2022 study, with significant variation in lipid content by trial month (Figure 3B; d.f. = 7, χ^2^ = 33.68, *p* < 0.0001).

The months of June (2.4%), September (2.3%), and January (2.3%) had the highest mean percent lipid, while November (1.8%) had the lowest. Significant differences in lipid content were observed between October and September (Steel-Dwass test: *z* =-3.08, *p* = 0.0433), and November and September (Steel-Dwass test: *z* =-3.64, *p* = 0.0067). Bee lipid content in June also differed significantly from bees sampled in July, August, October, and November (Table S4).

### 2.1 Seasonal variation in available pollen diet impacts the tolerance of bees to IAPV infection

To investigate whether seasonally dependent pollen nutrition affects honey bee tolerance to viral infection, we performed a follow-up study that tested whether the seasonality of available pollen forage, and, by extension, fluctuations in host nutritional status, was a contributing factor to our findings from the longitudinal tolerance experiment. We found that bees fed different seasonally dependent pollen diets (i.e. spring-collected vs fall-collected pollen) exhibited differences in their survivorship based on whether experimental-inoculation was performed on spring-emerged vs fall-emerged bees (Figure 4). Irrespective of diet treatment, spring-emerged bees that had been orally inoculated with IAPV exhibited significantly lower survivorship (69%) compared to non-inoculated controls (99%) after 72 hpi (Figure 4A; Kruskal-Wallis Test: d.f. = 1, χ^2^ = 60.68, *p* < 0.0001). A similar significant decline in survivorship was observed for fall-emerged bees (Figure 4D; Kruskal-Wallis Test: d.f. = 1, χ^2^ = 46.31, *p* < 0.0001) between those inoculated with IAPV (79%) and the non-inoculated control bees (99%), confirming successful infection occurred in both trials. Diet alone did not significantly affect survivorship in spring-emerged bees (Figure 4B; Kruskal-Wallis Test: d.f. = 2, χ^2^ = 0.1380, *p* = 0.9333). However, while neither significantly differed from the sucrose control, there was a significant effect of diet alone on the survivorship of fall-emerged bees between those fed the spring (96%) vs fall (87%) pollen diets (Figure 4E; Steel-Dwass test: *z* = 2.38, *p* = 0.0451).

**Figure 4:**
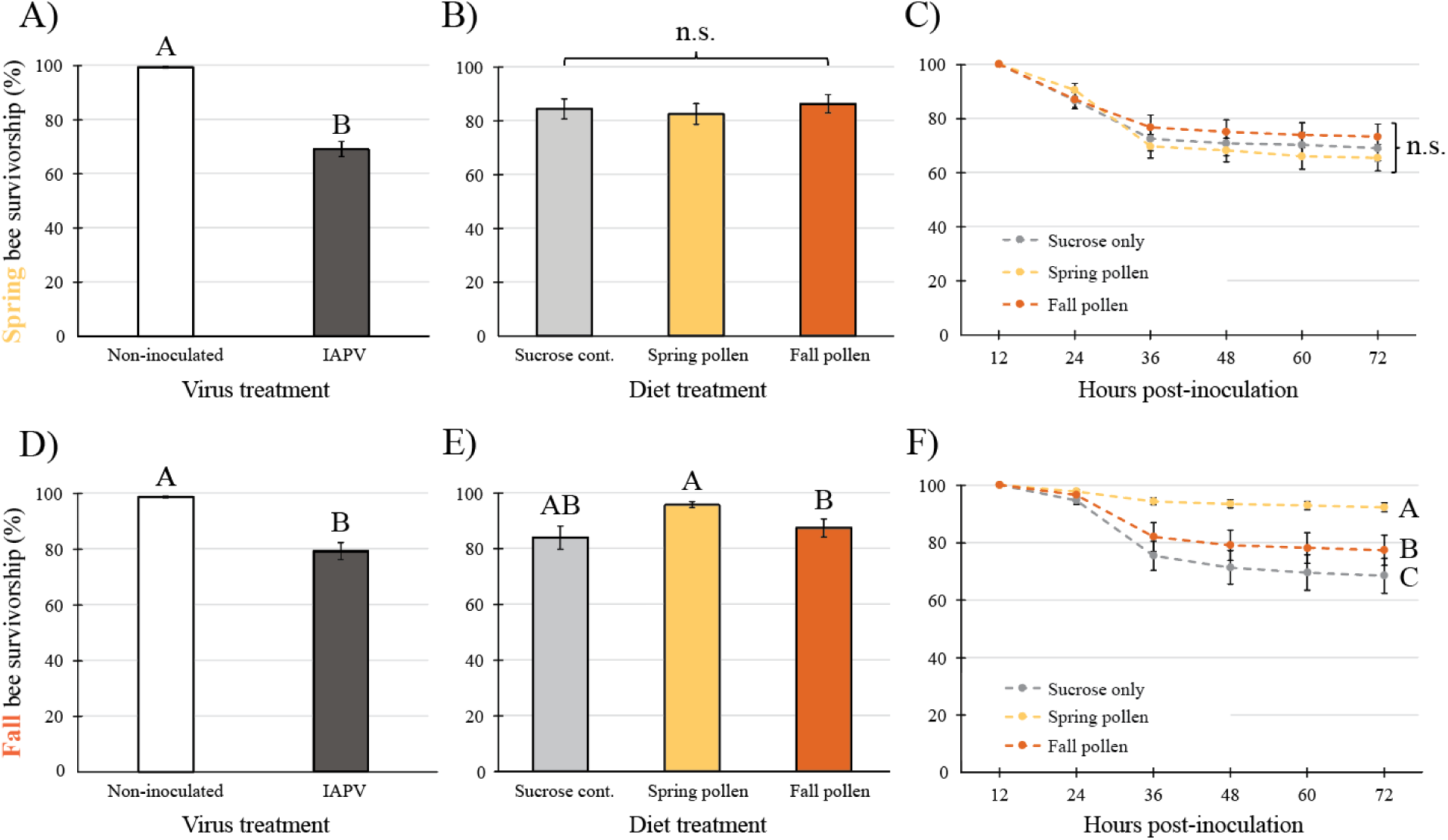
*Honey bee survivorship (%) after IAPV infection is influenced by seasonal variation in honey bee emergence and pollen diets.* Results are divided by honey bee seasonality (spring-emerged bees: panels A-C; fall-emerged bees: panels D-F). Bar graphs represent the mean (± s.e) cumulative proportion of surviving bees at 72 hpi, with significant differences signified by letters (a = 0.05). **A&D)** Comparisons of overall bee survivorship based on virus treatment alone (non-inoculated bees= white, IAPV-inoculated bees= gray). **B&E)** Comparisons of overall bee survivorship based on diet treatment alone. Caged bees were fed either spring-collected pollen (yellow), fall-collected pollen (orange), or a sucrose-only diet as a negative control (gray). **C&F)** Survivorship curves for IAPV-inoculated bees fed one of the different diet treatments (sucrose-only control= gray, spring-collected pollen= yellow, fall-collected pollen= orange).

Survivorship curves for the spring-emerged, IAPV-inoculated bees revealed no significance in survivorship between any of the three different diet treatments (Figure 4C). However, there was a significant effect of diet on the survivorship of fall-emerged bees, IAPV-inoculated bees (Figure 4F; d.f. = 2, χ^2^ = 91.28, *p* < 0.0001), with bees fed the spring pollen diet having significantly higher survivorship (92%) compared to bees fed the fall pollen (77%) or sucrose-only diets (68%). Fall-emerged bees inoculated with IAPV had a significantly higher risk of death when fed the fall pollen diet (Cox proportional hazards: HR= 2.96, *p* < 0.0001) or sucrose only control (Cox proportional hazards: HR= 4.18, *p* < 0.0001), compared to bees fed the spring pollen diet. Additionally, bees fed the sucrose-only diet had a higher risk of death compared to bees fed the fall pollen diet (Cox proportional hazards: HR= 1.41, *p* = 0.0033).

## 3. DISCUSSION

This study establishes that honey bees exhibit significant seasonal variation in their tolerance to viral infection, likely driven by time-dependent changes in host physiology and nutritional status throughout the year. Previous research has shown that seasonality affects certain pathogen-and environment-related disease factors known to influence the tolerance and susceptibility of honey bees to viral infection, including pathogen prevalence/abundance (7,9,50), *Varroa* abundance (7,9–11,17), and available forage within the environment (22,23). However, these studies do not address whether seasonality affects a honey bee’s ability to withstand viral infection.

Incidences of viral disease within honey bee colonies typically peak during the mid-summer to fall seasons, correlating with increased *Varroa* populations and the viruses they vector, as well as reduced forage availability, which can negatively affect bee nutrition and immune function (25,26). Our longitudinal tolerance experiment resulted in a similar seasonal trend, as honey bees experimentally infected with IAPV had the highest mortality and risk of death when inoculated during the months of August and September (Figure 1A-C). Conversely, bees inoculated during the summer (i.e., June and July) had the lowest overall mortality and risk of death. Quantifying the IAPV load of bees that had died as a result of infection during the longitudinal tolerance experiment demonstrated that a significantly lower mean IAPV quantity was associated with lethal infection in bees when they were inoculated in August-September compared to the other time periods (Figure 2). This was despite consistently low *Varroa* populations (Figure S1) and background viral loads (Table S3) within sampled colonies throughout the course of the study. In fact, background virus titers remained low during the month in which honey bees were observed to have the highest rates of mortality following IAPV inoculation (i.e., August). Thus, we concluded that bees inoculated with IAPV are less tolerant to viral infection in the months of August and September compared to bees inoculated during the rest of the year.

One explanation for this seasonal variation could be that, as a superorganism, honey bee colonies exhibit distinct seasonally dependent phenotypes, with significant physiological and behavioral differences occurring between summer and winter worker populations (51,52). In previous studies, located within a similar geographic location and climate as our own (37,53), the months of August and September were considered part of the pre-overwintering period (August-October) during which a colony will transition from a summer to winter population. This timeframe of winter bee production and population turnover has also been observed in more northern geographic regions (54). We hypothesize that the pre-overwintering period may be a time of increased vulnerability as the colony turns over its worker population and differentially allocates resources to better deal with more winter-related stressors, such as thermoregulation (31,36).

Multiple mechanisms, including honey bee nutritional state, have been proposed to control or contribute to this transition from summer to winter bees. For example, some studies tracking lipid content suggest that changes in this indicator of host nutritional status could be a driver in the transition between seasonal host phenotypes (reviewed in: Döke et al., 2015). Our studies measuring colony-level lipid content revealed significant seasonal fluctuations in lipid stores, with peak levels occurring in the months of June, September, and January (Figure 3).

These fluctuations closely align with our observations of IAPV mortality, particularly with high lipid content in June coinciding with our observed low mortality rates and a decline in lipid stores during the fall correlating with increased IAPV-induced mortality. While the lipid content did not perfectly align with our IAPV load data, it provides intriguing evidence linking honey bee nutrition and pathogen tolerance. Previous studies have shown that nutrient fluctuations may play a key role in honey bee health and immune function (55–57), further suggesting that nutritional deficits in the fall and winter months contribute to reduced viral tolerance.

In addition, honey bee colonies naturally reduce population growth in the fall as they prepare for the overwintering period. Short-lived summer bees are replaced by long-lived overwintering (i.e., diutinus) bees, which are characterized by higher nutritional stores (32). Following a peak in lipid stores in September, potentially driven by reduced forage and changes in regional pollen availability (58), new bees produced during this period are increasingly likely to exhibit diutinus traits, such as higher fat stores. However, at the onset of this period, diutinus bees comprise a smaller proportion of the colony population (51), meaning colony-level lipid stores remain low. As the year progresses and short-lived summer bees die off, the proportion of long-lived, fatty diutinus bees increases, leading to higher lipid stores at the colony level. These fluctuations suggest that honey bees differentially allocate energy for cellular function (59), and further highlight differences in immune function between summer and winter bees, with winter bees exhibiting reduced immune gene expression and heightened susceptibility to viral infections (60). This seasonal decline in lipid stores, particularly during the transition from summer to winter bees, gives further credence that this is a critical period of vulnerability, where reduced lipid content and altered immune function can contribute to decreased tolerance to viral infections. This underscores the tight link between honey bee nutritional status and immune function, which together influence viral susceptibility and tolerance.

Furthermore, our follow-up study investigating the effects of different seasonal pollen diets on viral tolerance supports the hypothesis that nutrition plays a critical role in honey bee tolerance to infection. While diet did not significantly affect viral tolerance in spring-emerged bees infected with IAPV (Figure 4C), fall-emerged bees showed the highest survivorship when fed a spring-collected pollen diet (Figure 4F). This aligns with previous studies, which found that honey bee responses to pathogens, such as *Nosema*, were influenced by the seasonal timing of available pollen (25). Other studies have similarly seen that spring-collected pollens, regardless of region, had higher amino acid and fatty acid concentrations compared to fall pollens (56), which may explain why fall-emerged bees fed spring pollen in our study responded better in terms of survivorship. These findings suggest that seasonal differences in pollen nutrition may modulate host immunity and pathogen tolerance, particularly during the fall transition. For the purpose of this study, we did not identify the different floral species within each seasonal pollen type. However, future studies aimed at better understanding the interactions between seasonality and honey bee nutrition, including the effects of different nutritional compositions on honey bee pathogen tolerance, could be used to develop methods in which to create honey bee diets aimed at enhancing winter bee resiliency to infection. This effect may be more pronounced in regions with critical gaps in high-quality fall forage, such as the Midwestern U.S. (58,61), which routinely reports high colony losses (14,16) and may benefit from additions of high-quality habitat that prioritize late season floral blooms.

Overall, our findings show that honey bee tolerance to viral infection varies significantly across the seasonal cycle, likely due to demographic shifts from summer to overwintering worker populations and the resulting fluctuations in colony nutritional status. This seasonality must be considered when evaluating disease risks and treatment strategies for honey bee colonies, such as interpreting longitudinal pathogen incidence and intensity data (e.g., Faurot-Daniels et al., 2020). For instance, understanding the times of heightened vulnerability, such as during the fall transition, can help optimize pathogen management strategies. Moreover, these insights are crucial for improving our ability to predict disease outbreaks and target effective treatments, particularly in the context of preparing colonies for overwintering. This research contributes to a deeper understanding of host-pathogen dynamics in honey bee colonies, emphasizing the need for seasonally tailored interventions to mitigate disease risks and enhance honey bee resilience. This is further emphasized by the recent, severe colony losses that has been attributed to honey bee viruses and their vectors (19). By integrating seasonal factors into disease models, beekeepers can better plan for times of high disease susceptibility, ultimately improving colony health and survival.

## 4. METHODS

### 4.1 Source and maintenance of honey bee colonies

Honey bees were sourced from research colonies managed by the University of Illinois Urbana-Champaign Bee Research Facility within Champaign County, Illinois. All colonies were headed by a queen of either Carniolan (*Apis mellifera carnica*) or Italian (*Apis mellifera ligustica*) descent and were maintained according to standard beekeeping practices, including regular *Varroa* mite monitoring and treatment (62,63). Research colonies were selected based on containing sufficient and approximately equal food stores, no overt signs of pathogen infection, and a low mite population (i.e., <3% threshold). Mite loads were determined monthly from June 2020-November 2020 using an alcohol wash method (64) and were continued into 2021 for select months (March, May, June, July, and October) for continuous confirmation of low mite populations while minimizing sampling impact on colony stability. From February 1 to April 14, 2021, and from November 16 to January 31, 2022, colonies were housed within a temperature-controlled (∼4°C) building to minimize cold stress during overwintering (65). Outside of the overwintering period, colonies were kept directly outside of the temperature-controlled building on the research facility grounds.

### 4.1 Creation of the IAPV inoculum

A purified IAPV inoculum was prepared for the longitudinal tolerance experiment following a previously described virus particle extraction and enrichment protocol (38). In short, white-eyed pupae were injected with a 1% solution of a previously purified IAPV inoculum and incubated under in-hive conditions (34°C and ∼50% relative humidity) to allow for IAPV propagation to occur. Virus particles were then extracted from the pupae through centrifugation and purified using polyethylene glycol (PEG) precipitation. The purity (>99% IAPV) and concentration (∼9.86 x 10^7^ g.e. of IAPV within 100 ng of purified total RNA) of the IAPV inoculum were determined via RNA extraction and RT-qPCR as described below (section 4.4). Due to its extremely high virulency, the inoculum was diluted 4-fold to create our high-end IAPV inoculum treatment (i.e., high dose; ∼9.86 x 10^3^ g.e. of IAPV within 100 ng of purified total RNA). Using the high dose as our baseline, we created five serial dilutions (i.e., 10^-2^, 10^-4^, 10^-6^, 10^-8^, 10^-10^) for our dose-response assays. Aliquots of each IAPV treatment were stored at-80°C for future experiments.

### 4.1 Establishment of the longitudinal tolerance experiment

The longitudinal tolerance experiment involved monthly collections of adult honey bee workers from research colonies, with colonies selected based on predefined criteria (section 4.1). To control for genetic variation, five research colonies were sampled consistently throughout the study from February 2021-January 2022 (n= 12 trial months). When colony-level mortality began in October 2021, additional colonies were incorporated to maintain a sample size of four to five colonies per month (Table S5). A subset of adult workers was randomly selected from the brood nest of each colony and placed into acrylic cube cages (10×10×8 cm; n= 5 bees/cage), ensuring standardized sampling. Each cage was equipped with a gravity sucrose feeder (30% sucrose, w:v) and a gravity water feeder, which were refilled *ad libitum*.

Cages were randomly assigned to our previously described IAPV inoculum treatment groups (n= 8 treatments), which consisted of a high dose of IAPV, its five dilutions (10^-2^, 10^-4^, 10^-6^, 10^-8^, 10^-10^), and two negative controls: one that controlled for stress-induced mortality due to the injection process itself (i.e., injection control) and one in which bees were not injected at all that controlled for any stressors attributed to the handling and anesthetization process (i.e., handling control). Within each trial month, each treatment was replicated with 4-5 cages. Bees were anesthetized with CO_2_ and then injected using a multi-dispenser pipette and 30G needle with 1 µL of their respective treatment, excluding the handling control, using standardized procedures (38,66). After injection, bees were returned to their cages and incubated at 34 and ∼50% relative humidity. Mortality was recorded every 12 hours from 12 to 48 hours post-inoculation (hpi), with only mortality occurring within this timeframe considered in the analysis based on IAPV’s known infection timeline (57,67). Dead bees were collected daily during each trial month and stored at-80°C for later IAPV load analysis.

### 4.1 Quantification of viral load

Absolute IAPV quantification from dead bees sampled during the longitudinal tolerance experiment was determined using RT-qPCR via the standard curve method. Only bees that died due to experimental infection were analyzed, as they exhibit comparable infection patterns as live-collected bees (67–69) and avoid complications related to cage depopulation. Given the large number of treatment groups (12 trial months; 8 IAPV treatments; 3 mortality checkpoints; and 4-5 cages per treatment), a subset of samples was processed. We selected bees from the 10^-2^ IAPV treatment group, as this dose had dead bees for each trial month, and then binned samples by season (spring: February and March (n=22 bees); summer: May and June (n=24); fall: August and September (n=24); and winter: November and December (n=24). This categorization and grouping of months by season were based on the seasonal designation of bees in previous studies conducted in a similar region and climate as this study (37,53). Additionally, the background virus levels (Table S3) of research colonies used in the trial months of June, August, and December were determined to spot-check for naturally high IAPV levels and the presence of other commonly detected honey bee-associated viruses, including deformed wing virus (DWV), black queen cell virus (BQCV), and sacbrood virus (SBV).

Total RNA extractions were performed on each bee sample, which consisted of individual, whole-bodied extractions using the Qiagen RNeasy Mini Kit (QIAGEN) and following the manufacturer’s protocol. The concentration and purity of each sample was measured on a Nanodrop 1000 spectrophotometer. Absolute quantification was conducted using the virus-specific primers and RT-qPCR protocol described in Carrillo-Tripp et al. (2016). In short, the *Power* SYBR™ Green RNA-to-C_T_™ *1-Step* Kit (Applied Biosystems) and a QuantStudio 6 real-time PCR instrument were used, with a standard curve generated using eight serial dilutions (1:10) of an in-vitro transcript containing virus-specific sequences for IAPV, DWV, BQCV, and SBV (67). Each well reaction contained 20 ng of total RNA, with samples performed in technical duplicates. Virus quantities and respective CT values were obtained using QuantStudio RealTime PCR Software (Applied Biosystems, version 1.3, Foster City, CA USA).

The limit of detection (LOD), based on repeated serial dilutions tests of the standard (70), was 4.92E+02 viral genome equivalents (g.e.) for IAPV and BQCV and was 4.92E+03 g.e. for DWV and SBV (Carrillo-Tripp et al., 2016). Any results falling below these detection thresholds were considered “undetected.” Background virus levels were quantified using a protocol similar to the one described above, with a few modifications. Specifically, virus levels were assessed at the colony level by using group extractions of 15 randomly selected honey bee workers per colony, and 200 ng of total RNA (as opposed to 20 ng) was used as the template for each RT-qPCR reaction.

### 4.1 Establishment of the longitudinal lipid content experiments

A parallel, nutrition-based longitudinal experiment measured colony-level lipid content (i.e., fat stores) alongside the longitudinal tolerance experiment using the same research colonies, sampling protocol, and timeline described in section 4.3. Due to the limited colony sample size in this initial study, a follow up experiment was conducted to verify trends using a greater colony sample size. This second study analyzed samples previously collected from June 2020 to January 2021 across seven apiaries in Champaign County, Illinois. Colony samples were obtained in June 2020 (n= 46 colonies), July (n= 68), August (n= 37), September (n= 40), October (n= 58), November (n= 30), December (n= 7), and January 2021 (n= 11), with the reduced colony numbers in December and January reflecting colonies entering the overwintering period. Lipid extraction and quantification for both lipid studies followed a modified protocol of Toth and Robinson (2005) as described in (65,72). For each colony and trial month across both studies, 15 nurse bees were homogenized in liquid nitrogen, and approximately 0.3g of homogenate was extracted using a 2:1 chloroform-to-methanol ratio over 12 hours. Lipid content was quantified via phosphor-vanillin spectrophotometric assays and calculated as mg lipid/mg bee mass (i.e., % lipid).

### 4.1 Establishment of the seasonal pollen experiment

In a follow up study, we determined how the seasonality of honey bees (spring-emerged and fall-emerged bees) and available nutrition (spring-collected and fall-collected pollen) interact to influence honey bee susceptibility to IAPV infection. Two multifactor cage trials were conducted in May (i.e., the spring bee trial) and August (i.e., the fall bee trial) of 2023. For both trials, cages (n= 35 bees/cage) were randomly assigned to a virus and diet treatment group. Virus treatment consisted of caged bees fed either: 1) 600 µL of a 10^-2.5^ µg/µL dose (∼3.12 x 10^5^ g.e. of IAPV within 100 ng of purified total RNA) of the previously described, undiluted IAPV stock mixed into 30% sucrose (w: v), or 2) 600µL of an unadulterated 30% sucrose solution that acted as a negative control (i.e., non-inoculated bees). Solutions were administered to cages via a weigh boat placed inside the cage, during which no other sucrose, water, or pollen was available. At 12 hpi, weigh boats were removed, and sucrose feeders containing unadulterated 50% sucrose solution were placed into each cage and refilled *ad libitum*. Additionally, cages were administered their respective diet treatment at 12 hpi, which consisted of either spring-collected pollen, fall-collected pollen, or a no pollen (i.e., sucrose-only) control. Experimental pollen was collected via two hive entrance pollen traps over a 48 hour-period in both April 2021 (i.e., spring-collected pollen) and August 2021 (i.e., fall-collected pollen). Hives for pollen collection were maintained at local apiaries managed by the Bee Research Facility. Collected pollen was homogenized by season and ground up via table top coffee grinder. Caged bees received their respective pollen diets via a modified 1.5 mL tube, which was refilled daily. In total, the study included twelve treatment groups (2 seasonal trials, 3 diet treatments, and 2 virus treatments), with 15 cage replicates per treatment group. Due to a high demand for bees in other spring experiments, the spring trial was set up over two days, with half of the cages set up each day. Mortality was recorded every 12 hours over a 72-hour period, and dead bees were removed at each checkpoint.

### 4.1 Statistical analysis

Analyses were performed with a combination of R version 4.3.2 (R Core Team, 2023) and JMP statistical software (JMP®, Pro 17. SAS Institute Inc., Cary, NC, 1989–2023). For each dataset, normality was assessed with quantile plots and the Shapiro-Wilk Goodness-of-Fit Test, and the level of significance was set at α = 0.05. For the longitudinal tolerance experiment, we conducted Chi-square tests, Cox proportional hazard models, and Effect Likelihood Ratio Tests. Chi-square tests determined the cumulative mortality at 48 hpi for each of the IAPV treatment groups within each trial month (Figure 1A; Table S1), with the injection control acting as the comparison group to which we compared all other treatments against within each respective month. Survival analyses using Cox proportional hazard models (Figure 1B) were performed via the survival package and visualized using the survminer package for the 10^-4^ IAPV treatment group over 48 hpi. Estimated hazard ratios of the 10^-4^ IAPV treatment group at 48 hpi for each trial month (Figure 1C; Table S2) were determined using Effect Likelihood Ratio Tests, with the month of August acting as the reference comparison group due to having the overall highest level of mortality. Pairwise comparisons incorporated Benjamini-Hochberg corrections to reduce Type 1 errors. Virus quantities obtained via RT-qPCR were first log-transformed and then binned by season (see section 4.4). After failing to meet normality assumptions, significance was determined using Kruskal-Wallis rank sum tests and post hoc pairwise comparisons using Wilcoxon rank sum tests with Benjamini-Hochberg corrections (Figure 2). For the parallel 2021-2022 lipid study, significance in mean % lipid values between months (Figure 3A) was determined using a one-way ANOVA, followed by a Tukey HSD post hoc test, with colony ID included as a random effect. For the larger-scale 2020-2021 lipid study (Figure 3B; Table S4), a Kruskal-Wallis Test was performed followed by a post hoc Steel-Dwass test. For the seasonal pollen experiment, significant differences in endpoint mortality based on the virus and diet treatments alone (Figure 4A&D and 4B&E, respectively) were determined via Kruskal-Wallis rank sum tests, followed by post hoc pairwise comparisons using Wilcoxon rank sum tests with Benjamini-Hochberg corrections. Survival analyses (4C and 4F) used Cox proportional hazard models were used to determine the association between diet treatment and honey bee seasonality on the survival outcome of IAPV-inoculated bees, with cage as a random effect within the models. Penalized log-likelihood estimation was used to regularize the models.

## ACKNOWLEDGEMENTS

We would like to thank Nathan Beach for his valuable contributions as apiary manager in maintaining research colonies and to Hannah Salzburg for helping with the seasonal pollen experiment.

## AUTHOR CONTRIBUTIONS

Conceptualization: AGD, ALS, GPH; Investigation: ALS, ANP, GPH, LNT, MS, VP; Formal Analysis: ALS, ANP, GPH; Data Curation: ALS, ANP, GPH, VP; Writing—Original Draft Preparation: ANP; Writing—Review and Editing: AGD; ALS, ANP; Supervision: AGD; Project Administration: AGD; Funding Acquisition: AGD.

## SUPPLEMENTARY TABLES & FIGURES

**Table S1:** All pairwise Chi-square tests determining statistical differences in cumulative mortality at 48 hpi for the longitudinal tolerance experiment. Within each month, the average proportion of mortality for the injection control group was compared against each of the other IAPV treatments (i.e., the high dose, its five dilutions, and the handling control) to test for significance. Resulting p values were adjusted using the Benjamini-Hochberg method. Asterisks under the “Sig.” column indicate significant comparisons.

**Table S2:** Estimated hazard ratios of the longitudinal tolerance experiment, generated from Effect Likelihood Ratio Tests and Benjamini-Hochberg corrections. Comparisons were made for the 10^-4^ IAPV treatment group at 48 hpi between each trial month and the month of August.

August acted as the reference comparison group due to having the overall highest level of mortality. Asterisks under the “Sig.” column indicate significant comparisons.

**Table S3:** Background viral loads of research colonies used for the longitudinal tolerance experiment. Background levels include the CT means and genome equivalent quantities of common honey bee-infecting viruses, including deformed wing virus (DWV), black queen cell virus (BQCV), sacbrood virus (SBV), and our virus of interest: Israeli acute paralysis virus (IAPV), for each of the 4-5 research colonies from which bees were sampled from for the months of June, August, and December.

**Table S4:** All pairwise comparisons of the larger-scale 2020-2021 longitudinal lipid content study comparing the mean % lipid content of colonies between months using a Kruskal-Wallis Test followed by a post hoc Steel-Dwass test. Asterisks under the “Sig.” column indicate significant comparisons.

**Table S5:** Timeline of colonies used for the longitudinal tolerance experiment and parallel lipid study. Honey bee workers were sampled from the same colonies from February through

September. In October, colonies with low *Varroa* levels were selected to replace those that had failed.

**Figure S1:** Mean *Varroa* mite counts of colonies used for the longitudinal tolerance experiment. Values depict the mean mite count (per 300 bees) of all colonies sampled at each monthly timepoint. In 2020, mean mite counts were averaged from ∼65 colonies across 10 local apiaries from June-November. Colonies with the lowest mite counts were selected for the longitudinal tolerance experiment. Mite sampling of our experimental colonies continued throughout the course of the experiment. However, mite counts were not performed during winter months while colonies were in cold storage, in order to reduce stress and maintain the smaller worker populations throughout the overwintering period. The red dotted line represents the 3% economic threshold, above which *Varroa* infestation levels are commonly considered high and at risk of negatively impacting colony health.

## REFERENCES

1. Altizer S, Dobson A, Hosseini P, Hudson P, Pascual M, Rohani P. Seasonality and the dynamics of infectious diseases. Ecology Letters. 2006;9(4):467–84.

2. Dowell SF. Seasonal variation in host susceptibility and cycles of certain infectious diseases. Emerg Infect Dis. 2001;7(3):369–74.

3. Fisman DN. Seasonality of Infectious Diseases. Annual Review of Public Health. 2007 Apr 21;28(Volume 28, 2007):127–43.

4. Grassly NC, Fraser C. Seasonal infectious disease epidemiology. Proceedings of the Royal Society B: Biological Sciences. 2006 Jul 7;273(1600):2541–50.

5. McNew GL. The nature, origin and evolution of parasitism. In: Horsfall, J G; Dimond, AE (Ed) Plant pathology: an advanced treatise New York, Academic Press. 1960;

6. Scholthof KBG. The disease triangle: pathogens, the environment and society. Nat Rev Microbiol. 2007 Feb;5(2):152–6.

7. Faurot-Daniels C, Glenny W, Daughenbaugh KF, McMenamin AJ, Burkle LA, Flenniken ML. Longitudinal monitoring of honey bee colonies reveals dynamic nature of virus abundance and indicates a negative impact of Lake Sinai virus 2 on colony health. PLOS ONE. 2020 Sep 8;15(9):e0237544.

8. Grozinger CM, Flenniken ML. Bee Viruses: Ecology, Pathogenicity, and Impacts. Annual Review of Entomology. 2019;64(1):205–26.

9. Runckel C, Flenniken ML, Engel JC, Ruby JG, Ganem D, Andino R, et al. Temporal Analysis of the Honey Bee Microbiome Reveals Four Novel Viruses and Seasonal Prevalence of Known Viruses, Nosema, and Crithidia. PLOS ONE. 2011 Jun 7;6(6):e20656.

10. Genersch E, Aubert M. Emerging and re-emerging viruses of the honey bee (Apis mellifera L.). Vet Res. 2010;41(6):54.

11. Traynor KS, Mondet F, Miranda JR de, Techer M, Kowallik V, Oddie MAY, et al. Varroa destructor: A Complex Parasite, Crippling Honey Bees Worldwide. Trends in Parasitology. 2020 Jul 1;36(7):592–606.

12. Francis RM, Nielsen SL, Kryger P. Varroa-Virus Interaction in Collapsing Honey Bee Colonies. PLOS ONE. 2013 Mar 19;8(3):e57540.

13. Sumpter DJT, Martin SJ. The dynamics of virus epidemics in Varroa-infested honey bee colonies. Journal of Animal Ecology. 2004;73(1):51–63.

14. Aurell D, Bruckner S, Wilson M, Steinhauer N, Williams GR. A national survey of managed honey bee colony losses in the USA: Results from the Bee Informed Partnership for 2020– 21 and 2021–22. Journal of Apicultural Research. 2024 Jan 1;63(1):1–14.

15. Berthoud H, Imdorf A, Haueter M, Radloff S, Neumann P. Virus infections and winter losses of honey bee colonies (Apis mellifera). Journal of Apicultural Research. 2010 Jan 1;49(1):60–5.

16. Bruckner S, Wilson M, Aurell D, Rennich K, vanEngelsdorp D, Steinhauer N, et al. A national survey of managed honey bee colony losses in the USA: results from the Bee Informed Partnership for 2017–18, 2018–19, and 2019–20. Journal of Apicultural Research. 2023 May 27;62(3):429–43.

17. Dainat B, Evans JD, Chen YP, Gauthier L, Neumann P. Predictive Markers of Honey Bee Colony Collapse. PLOS ONE. 2012 Feb 23;7(2):e32151.

18. Steinhauer N, Aurell D, Bruckner S, Wilson M, Rennich K, vanEngelsdorp D, et al. United States honey bee colony losses 2022–23: Preliminary results from the Bee Informed Partnership. JournalTitle Bee Informed [Internet]. 2023 [cited 2024 Mar 31]; Available from: https://beeinformed.org/2023/06/22/united-states-honey-bee-colony-losses-2022-23-preliminary-results-from-the-bee-informed-partnership/

19. Lamas ZS, Rinkevich F, Garavito A, Shaulis A, Boncristiani D, Hill E, et al. Viruses and vectors tied to honey bee colony losses [Internet]. bioRxiv; 2025 [cited 2025 Jun 6]. p. 2025.05.28.656706. Available from: https://www.biorxiv.org/content/10.1101/2025.05.28.656706v1

20. Medzhitov R, Schneider DS, Soares MP. Disease Tolerance as a Defense Strategy. Science. 2012 Feb 24;335(6071):936–41.

21. Lau P, Bryant V, Ellis JD, Huang ZY, Sullivan J, Schmehl DR, et al. Seasonal variation of pollen collected by honey bees (Apis mellifera) in developed areas across four regions in the United States. PLOS ONE. 2019 Jun 12;14(6):e0217294.

22. Synge AD. Pollen Collection by Honeybees (Apis mellifera). Journal of Animal Ecology. 1947;16(2):122–38.

23. Rodriguez Messan M, Page RE, Kang Y. Effects of vitellogenin in age polyethism and population dynamics of honeybees. Ecological Modelling. 2018 Nov 24;388:88–107.

24. Smart M, Pettis J, Rice N, Browning Z, Spivak M. Linking Measures of Colony and Individual Honey Bee Health to Survival among Apiaries Exposed to Varying Agricultural Land Use. PLOS ONE. 2016 Mar 30;11(3):e0152685.

25. DeGrandi-Hoffman G, Gage SL, Corby-Harris V, Carroll M, Chambers M, Graham H, et al. Connecting the nutrient composition of seasonal pollens with changing nutritional needs of honey bee (*Apis mellifera* L.) colonies. Journal of Insect Physiology. 2018 Aug 1;109:114– 24.

26. Dolezal AG, Toth AL. Feedbacks between nutrition and disease in honey bee health. Current Opinion in Insect Science. 2018 Apr 1;26:114–9.

27. Casadevall A, Pirofski L anne. What Is a Host? Attributes of Individual Susceptibility. Infection and Immunity. 2018 Jan 22;86(2):10.1128/iai.00636-17.

28. Van Sluijs L, Pijlman GP, Kammenga JE. Why do Individuals Differ in Viral Susceptibility? A Story Told by Model Organisms. Viruses. 2017 Oct;9(10):284.

29. Bresnahan ST, Döke MA, Giray T, Grozinger CM. Tissue-specific transcriptional patterns underlie seasonal phenotypes in honey bees (Apis mellifera). Molecular Ecology. 2022;31(1):174–84.

30. Dostálková S, Dobeš P, Kunc M, Hurychová J, Škrabišová M, Petřivalský M, et al. Winter honeybee (Apis mellifera) populations show greater potential to induce immune responses than summer populations after immune stimuli. Journal of Experimental Biology. 2021 Feb 8;224(3):jeb232595.

31. Fahrenholz L, Lamprecht I, Schricker B. Thermal investigations of a honey bee colony: thermoregulation of the hive during summer and winter and heat production of members of different bee castes. J Comp Physiol B. 1989 Sep 1;159(5):551–60.

32. Fluri P, Lüscher M, Wille H, Gerig L. Changes in weight of the pharyngeal gland and haemolymph titres of juvenile hormone, protein and vitellogenin in worker honey bees. Journal of Insect Physiology. 1982 Jan 1;28(1):61–8.

33. Kešnerová L, Emery O, Troilo M, Liberti J, Erkosar B, Engel P. Gut microbiota structure differs between honeybees in winter and summer. The ISME Journal. 2020 Mar 1;14(3):801–14.

34. Lee S, Kalcic F, Duarte IF, Titera D, Kamler M, Mrna P, et al. 1H NMR Profiling of Honey Bee Bodies Revealed Metabolic Differences between Summer and Winter Bees. Insects. 2022 Feb;13(2):193.

35. Maurizio A, Hodges FED. The Influence of Pollen Feeding and Brood Rearing on the Length of Life and Physiological Condition of the Honeybee Preliminary Report. Bee World. 1950 Feb;31(2):9–12.

36. Stabentheiner A, Pressl H, Papst T, Hrassnigg N, Crailsheim K. Endothermic heat production in honeybee winter clusters. Journal of Experimental Biology. 2003 Jan 15;206(2):353–8.

37. Ward R, Coffey M, Kavanagh K. Proteomic characterisation of the summer–winter transition in Apis mellifera. Apidologie. 2022 Aug 2;53(4):39.

38. Hsieh EM, Carrillo-Tripp J, Dolezal AG. Preparation of Virus-Enriched Inoculum for Oral Infection of Honey Bees (Apis mellifera). JoVE (Journal of Visualized Experiments). 2020 Aug 26;(162):e61725.

39. Chen YP, Pettis JS, Corona M, Chen WP, Li CJ, Spivak M, et al. Israeli Acute Paralysis Virus: Epidemiology, Pathogenesis and Implications for Honey Bee Health. PLOS Pathogens. 2014 Jul 31;10(7):e1004261.

40. Amiri E, Seddon G, Zuluaga Smith W, Strand MK, Tarpy DR, Rueppell O. Israeli Acute Paralysis Virus: Honey Bee Queen–Worker Interaction and Potential Virus Transmission Pathways. Insects. 2019 Jan;10(1):9.

41. Boncristiani HF, Evans JD, Chen Y, Pettis J, Murphy C, Lopez DL, et al. In Vitro Infection of Pupae with Israeli Acute Paralysis Virus Suggests Disturbance of Transcriptional Homeostasis in Honey Bees (Apis mellifera). PLOS ONE. 2013 Sep 5;8(9):e73429.

42. Geffre AC, Gernat T, Harwood GP, Jones BM, Morselli Gysi D, Hamilton AR, et al. Honey bee virus causes context-dependent changes in host social behavior. Proceedings of the National Academy of Sciences. 2020 May 12;117(19):10406–13.

43. McCormick EC, Cohen OR, Dolezal AG, Sadd BM. Consequences of microsporidian prior exposure for virus infection outcomes and bumble bee host health. Oecologia. 2023 Jun 1;202(2):325–35.

44. Taylor LN, Dolezal AG. The effect of Israeli acute paralysis virus infection on honey bee brood care behavior. Sci Rep. 2024 Jan 10;14(1):991.

45. Dolezal AG, Hendrix SD, Scavo NA, Carrillo-Tripp J, Harris MA, Wheelock MJ, et al. Honey Bee Viruses in Wild Bees: Viral Prevalence, Loads, and Experimental Inoculation. PLOS ONE. 2016 Nov 10;11(11):e0166190.

46. Harwood GP, Prayugo V, Dolezal AG. Butenolide Insecticide Flupyradifurone Affects Honey Bee Worker Antiviral Immunity and Survival. Frontiers in Insect Science [Internet]. 2022 [cited 2024 Feb 5];2. Available from: https://www.frontiersin.org/articles/10.3389/finsc.2022.907555

47. Delaplane K S, Hood W Michael. Economic threshold for Varroa jacobsoni Oud. in the southeastern USA. Apidologie. 1999;30(5):383–95.

48. Gisder S, Aumeier P, Genersch E. Deformed wing virus: replication and viral load in mites (Varroa destructor). Journal of General Virology. 2009;90(2):463–7.

49. Kevill JL, Highfield A, Mordecai GJ, Martin SJ, Schroeder DC. ABC Assay: Method Development and Application to Quantify the Role of Three DWV Master Variants in Overwinter Colony Losses of European Honey Bees. Viruses. 2017 Nov;9(11):314.

50. D’Alvise P, Seeburger V, Gihring K, Kieboom M, Hasselmann M. Seasonal dynamics and co-occurrence patterns of honey bee pathogens revealed by high-throughput RT-qPCR analysis. Ecology and Evolution. 2019;9(18):10241–52.

51. Döke MA, Frazier M, Grozinger CM. Overwintering honey bees: biology and management. Current Opinion in Insect Science. 2015 Aug 1;10:185–93.

52. Winston ML. The Biology of the Honey Bee. Harvard University Press; 1991. 300 p.

53. Quinlan GM, Grozinger CM. Evaluating the role of social context and environmental factors in mediating overwintering physiology in honey bees (Apis mellifera). Journal of Experimental Biology. 2024 Mar 22;jeb.247314.

54. Mattila HR, Harris JL, Otis GW. Timing of production of winter bees in honey bee (Apis mellifera) colonies. Insectes soc. 2001 Jun 1;48(2):88–93.

55. Alaux C, Ducloz F, Crauser D, Le Conte Y. Diet effects on honeybee immunocompetence. Biology Letters. 2010 Jan 20;6(4):562–5.

56. DeGrandi-Hoffman G, Corby-Harris V, Carroll M, Toth AL, Gage S, Watkins deJong E, et al. The Importance of Time and Place: Nutrient Composition and Utilization of Seasonal Pollens by European Honey Bees (Apis mellifera L.). Insects. 2021 Mar;12(3):235.

57. Dolezal AG, Carrillo-Tripp J, Judd TM, Allen Miller W, Bonning BC, Toth AL. Interacting stressors matter: diet quality and virus infection in honeybee health. Royal Society Open Science. 2019 Feb 6;6(2):181803.

58. Dolezal AG, St. Clair AL, Zhang G, Toth AL, O’Neal ME. Native habitat mitigates feast– famine conditions faced by honey bees in an agricultural landscape. Proceedings of the National Academy of Sciences. 2019 Dec 10;116(50):25147–55.

59. McKinstry M, Chung C, Truong H, Johnston BA, Snow JW. The heat shock response and humoral immune response are mutually antagonistic in honey bees. Sci Rep. 2017 Aug 18;7(1):8850.

60. Steinmann N, Corona M, Neumann P, Dainat B. Overwintering Is Associated with Reduced Expression of Immune Genes and Higher Susceptibility to Virus Infection in Honey Bees. PLOS ONE. 2015 Jun 29;10(6):e0129956.

61. Zhang G, St. Clair AL, Dolezal A, Toth AL, O’Neal M. Honey Bee (Hymenoptera: Apidea) Pollen Forage in a Highly Cultivated Agroecosystem: Limited Diet Diversity and Its Relationship to Virus Resistance. Journal of Economic Entomology. 2020 Jun 6;113(3):1062–72.

62. Currie RW, Gatien P. Timing acaricide treatments to prevent Varroa destructor (Acari: Varroidae) from causing economic damage to honey bee colonies. The Canadian Entomologist. 2006 Apr;138(2):238–52.

63. Gregorc A, Sampson B. Diagnosis of Varroa Mite (Varroa destructor) and Sustainable Control in Honey Bee (Apis mellifera) Colonies—A Review. Diversity. 2019 Dec;11(12):243.

64. Lee KV, Moon RD, Burkness EC, Hutchison WD, Spivak M. Practical Sampling Plans for Varroa destructor (Acari: Varroidae) in Apis mellifera (Hymenoptera: Apidae) Colonies and Apiaries. Journal of Economic Entomology. 2010 Aug 1;103(4):1039–50.

65. St. Clair AL, Beach NJ, Dolezal AG. Honey bee hive covers reduce food consumption and colony mortality during overwintering. PLOS ONE. 2022 Apr 4;17(4):e0266219.

66. Bailey L, Gibbs AJ, Woods RD. Two viruses from adult honey bees (*Apis mellifera* Linnaeus). Virology. 1963 Nov 1;21(3):390–5.

67. Carrillo-Tripp J, Dolezal AG, Goblirsch MJ, Miller WA, Toth AL, Bonning BC. In vivo and in vitro infection dynamics of honey bee viruses. Sci Rep. 2016 Feb 29;6(1):22265.

68. Hsieh EM, Berenbaum MR, Dolezal AG. Ameliorative Effects of Phytochemical Ingestion on Viral Infection in Honey Bees. Insects. 2020 Oct;11(10):698.

69. Payne AN, Prayugo V, Dolezal AG. A honey bee-associated virus remains infectious and quantifiable in postmortem hosts. Journal of Invertebrate Pathology. 2025 Mar 1;209:108258.

70. de Miranda JR, Bailey L, Ball BV, Blanchard P, Budge GE, Chejanovsky N, et al. Standard methods for virus research in Apis mellifera. Journal of Apicultural Research. 2013 Jan 1;52(4):1–56.

71. Toth AL, Robinson GE. Worker nutrition and division of labour in honeybees. Animal Behaviour. 2005 Feb 1;69(2):427–35.

72. Dolezal AG, Carrillo-Tripp J, Miller WA, Bonning BC, Toth AL. Intensively Cultivated Landscape and Varroa Mite Infestation Are Associated with Reduced Honey Bee Nutritional State. PLOS ONE. 2016 Apr 12;11(4):e0153531.

73. R Core Team [Internet]. 2023 [cited 2025 Jun 6]. Available from: https://www.r-project.org/

